# Disentangling shape and size variation to compare canopy architecture in maize

**DOI:** 10.64898/2026.09.17.752132

**Authors:** Kristian Johnson, Christian Fournier, Carine Palaffre, Randall J. Wisser

**Affiliations:** LEPSE, INRAE, L’Institut Agro, University of Montpellier, Montpellier 34060, France; Unité Expérimentale du Maïs, INRAE, Saint Martin De Hinx 40390, FR; Department of Plant and Soil Sciences, University of Delaware, Newark, DE, 19716 USA

**Keywords:** plant architecture, ontogeny, reproductive transition, flowering time, functional data analysis, Poaceae

## Abstract

Comparing maize canopies is confounded by two problems: plants differ in the number of phytomers they produce, and phytomer size scales with that number. Position along the shoot is therefore an unreliable basis for comparison, confounding shape with size.
We first position the floral transition (tassel initiation, T.I.) within shoot development, revealing it as the landmark that organizes a conserved canopy shape. This provides the foundation for aligning canopies that differ in phytomer number.
Focusing on the leaf blade, we used functional data analysis on a maize panel across controlled and field environments to decompose variation in blade-length profiles, which was dominated (77%) by overall amplitude, identifying scaling as the major source of size variation. Because blades enlarge with each phytomer until the transition, later T.I. yields a larger maximum; this T.I.-linked component explains up to one-third of scaling variation, reinforced by a shared genetic association between T.I. and blade size with the flowering-time gene *ZmCCT10*.
Regressing out this component leaves a residual that defines a new, heritable axis of intrinsic scaling, independent of flowering time. It distinguishes hyper- and hypo-scaling genotypes consistently across environments, offering a selection target decoupled from phenology.

**Plain Language Summary:** Plants vary in how many leaves they grow and how big those leaves get, making canopies hard to compare across different varieties and growing conditions. Studying maize, we found that the timing of flowering organizes the overall shape of the canopy, letting us compare them fairly, and uncovered a separate, inherited source of leaf-size variation. This could help breeders adjust crop canopies without changing when plants flower.

## Introduction

Plant canopy architecture is the three-dimensional spatial arrangement of the aboveground shoot system, governing how plants interact with the atmosphere through light interception, gas exchange, and water loss, which together drive growth and yield (Durand & Robson, 2023; Murchie & Burgess, 2022; Perez et al., 2019). It emerges from two developmental processes: organogenesis, the initiation and development of new organs, and extension, the growth of existing structures (Barthélémy & Caraglio, 2007). These processes converge at the level of the vegetative phytomer, the functional unit of the shoot system, whose development gives rise to architectural traits including plant height and the size, angle, and orientation of leaves (McAllister et al., 2025; Strable, 2021). Thus, understanding the coupling between phytomer development and its emergent morphology has broad implications in plant science, ranging from crop breeding and computational plant modeling to predicting how genotypes respond to different environments and how species adapt ecologically.

The vegetative phytomer is an individual unit of the shoot system, comprising a leaf attached to a node, an axillary bud, and an internode (McMaster, 2005). Monocots such as maize (*Zea mays* ssp. *mays* L.; focus of this study) are distinguished by a leaf with a blade and a sheath component that encircles the stem, which determines how successive phytomers overlap and develop through their repeated stacking along the stem axis (Conklin et al., 2019). The blade, sheath, and internode are therefore the basic components of canopy dimensioning and architecture. As they form sequentially within a plant, differences in their timing of initiation and duration of elongation produce characteristic gradients in size along the shoot that rise and fall across successive ranks. Representing the size gradient as a continuous function has shown that each component in maize forms a peaked profile with differences in amplitude and skewness: blades (Dwyer et al., 1992; Fan et al., 2020; Lacube et al., 2020; Vidal et al., 2021), sheaths (Andrieu et al., 2006; Vidal et al., 2021), and internodes (Birch et al., 2002; Fournier, 2000). However, how the sizes of these components change with respect to each other across time, and how profiles vary across genotypes and environments, remains largely unexplored.

Studies measuring variation in the shoot system often use either single-point measurements at specific positions (e.g., leaf angle or length at a given leaf rank) or aggregate measures (e.g., total leaf area) that collapse the canopy into a single value. Understanding the shoot as a complete structure requires a different approach, one in which the profiles of architectural dimensions are compared across phytomer positions. While the profile of each phytomer component exhibits a conserved structure, comparing these across plants presents fundamental challenges of both positional alignment and dimensional scaling (**Fig. 1**). First, plants vary in their final number of phytomers, making it unclear how to align their profiles or what a common rank position means. Second, because phytomer dimensions are known to scale with phytomer number (Ducrocq & Duru, 2000), as demonstrated for leaf-blade profiles of maize (Lacube et al., 2020), comparisons of size at common ranks will be confounded with development. The combined effects of alignment and scaling in canopy comparison has not been addressed, despite both being key bottlenecks to dissecting variation in canopy architecture and broader examples of how ontogenetic variation confounds biological comparison.

**Figure 1.**
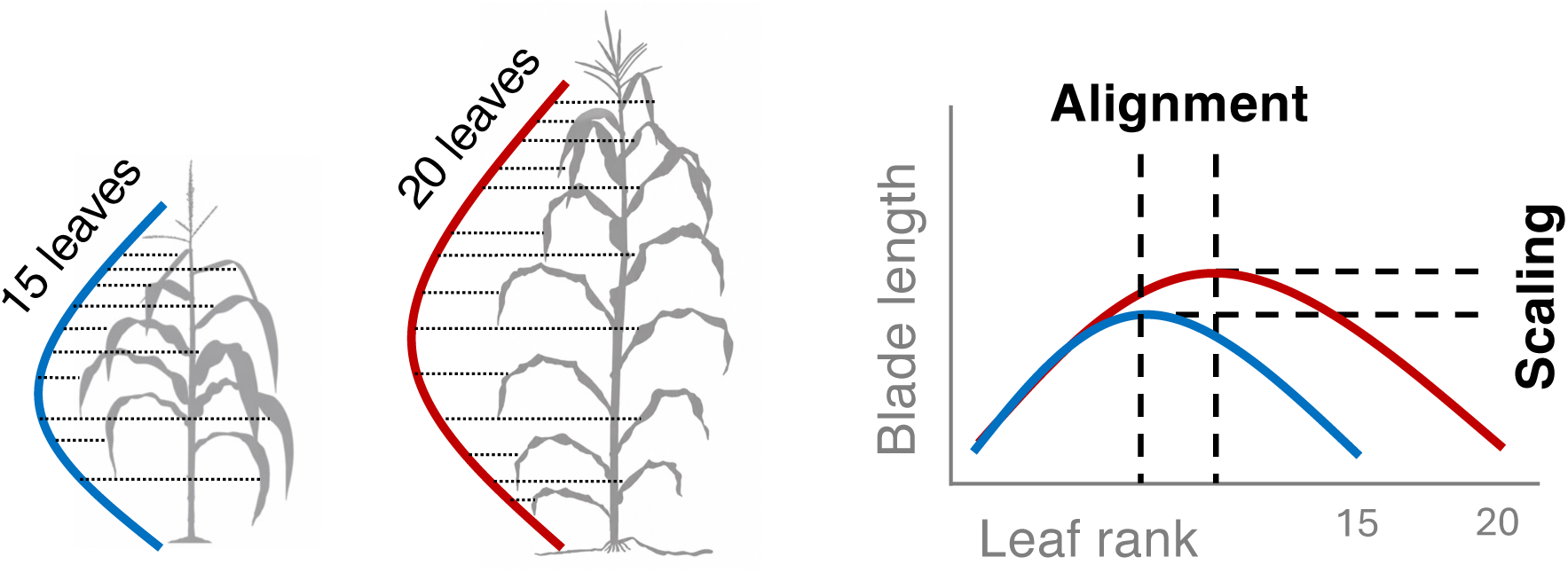
The dual problem of alignment and scaling in canopy architecture comparisons. This example illustrates how comparing blade-length profiles by absolute rank confounds two distinct effects. First, because profile shape is conserved in form, plants with different numbers of phytomers have profiles shifted along the rank axis that do not align (the alignment problem). Second, profile magnitude also varies with phytomer number, so these plants differ in profile height independently of shape (the scaling problem). On an absolute-rank axis these effects compound, preventing meaningful comparison; disentangling them is required to compare profiles on a common basis.

The alignment problem arises due to differences in final number of phytomers, which in maize is set by the timing of floral transition, or tassel initiation (T.I.). At T.I., the shoot apical meristem (SAM) ceases to produce vegetative phytomers and transforms into a reproductive meristem, forming the male flower (tassel) from the SAM and female flower (ear) from an axillary meristem (Colasanti & Muszynski, 2009; McSteen & Leyser, 2005). Variation in T.I. timing, and thus in final phytomer number, arises from both genetic and environmental effects (Drouault et al., 2025). Within a common field environment, genotypic differences in T.I. can cause final phytomer number to vary widely, from 13 to 31 leaves (this study). The environmental effect on leaf number is driven primarily by photoperiod, through its effect on T.I. timing (Kiniry et al., 1983). As a facultative short-day species, maize delays T.I. in photoperiod-sensitive genotypes under long daylengths, producing additional leaves relative to its basic leaf number in short-day conditions (Bonhomme et al., 1991; Ellis et al., 1992).

The scaling problem also likely stems from T.I., an association described decades ago (Borrill, 1959) that is now grounded in a model for the coordinated development of the maize shoot system (Fournier et al., 2005). During the initial phase of vegetative growth, each emerging leaf promotes development of the next, creating positive feedback that results in progressively larger organs until the floral transition (Andrieu et al., 2006; Casey et al., 1999; Skinner & Nelson, 1995). This pre-T.I. coordination operates through two developmental events—leaf tip and collar appearance—that regulate cellular division and expansion (Fournier et al., 2005; Skinner & Nelson, 1995). Tip emergence of each leaf initiates the formation of a collar, a boundary layer demarcating the blade and sheath, which also stabilizes blade elongation. As the collar emerges, it signals cessation of elongation in both the sheath and associated internode (Fournier, 2000; Vidal & Andrieu, 2020; Zhu et al., 2014). Before T.I., each successive phytomer has a longer path length to traverse for tip appearance, so phytomer size scales with phytomer number (Barre et al., 2015; Skinner & Nelson, 1995). At T.I., this coordination system shifts to a new dynamic where the blade-sheath boundary is formed at an increased rate independent of tip appearance, resulting in sequentially smaller sized blades and similarly sized sheaths (Andrieu et al., 2006; Vidal et al., 2021). Meanwhile, the release of apical dominance at T.I. (Martin et al., 1988; Zhang et al., 2019) leads to a secondary phase of rapid internode elongation (Morrison et al., 1994) and development of the dominant ear (in maize) from the final axillary bud (Baba & Yamazaki, 1996; Lejeune & Bernier, 1996).

Given that T.I. affects both final phytomer number and, as argued above, the developmental scaling of the shoot, we tested the hypothesis that T.I. determines the conserved shape and size-scaling relationship of phytomer profiles. We also questioned how to resolve the alignment and scaling problems for canopy comparison across genotypes and environments. To examine this at population scale, normally precluded by the serial dissections required to determine T.I. timing, we first improved the prediction of T.I. through a meta-analysis, allowing it to be positioned as a landmark within the profiles of phytomer components across individuals. We initially examined the blade, sheath, and internode components together to characterize how their profile shapes are structured around T.I. and coordinated through development. We then focused on the leaf blade, measuring its canopy-level length profiles in field experiments with diverse genotypes and contrasting photoperiod environments, and used functional data analysis (Yao et al., 2005) to compare registration systems for profile alignment and examine the effect of T.I. on scaling variation. From this, we present a standardized approach for whole-plant canopy comparison and identify a new axis of variation relevant to the genetics and breeding of maize canopy architecture.

## Materials and Methods

### Meta-analysis and formalism comparisons

Formalisms for predicting the timing of tassel initiation (T.I) or visible leaf tip at T.I include parameters for the plastochron (thermal time between the initiation of successive leaf primordia) to phyllochron (thermal time between the appearance of successive leaf tips) ratio (*α_plast:pℎyll_*) and the number of leaf primordia present at crop emergence, when the seedling is first visible above the soil (*L_em_*) (**Eq. 1** below). To obtain a robust estimate of *α_plast:pℎyll_*, a meta-analysis was performed using 11 of 14 studies reporting on the relation between the number of leaf primordia and leaf tips formed across time (**Table S1**). Excluded studies either measured collar appearance instead of leaf tips or used a contrasting definition of primordium size to describe an initiated leaf. With the exception of one study for which the original data were provided by the authors (Padilla & Otegui, 2005), data for the remaining studies was extracted from figures using DigitizeIt (Braunschweig, Germany); i.e., data points in each study figure were converted to numerical data for statistical analysis. The *α_plast:pℎyll_* was estimated per study as the slope coefficient obtained by simple linear regression of the number of leaf primordia as the response variable and the number of leaf tips as the predictor variable. For studies containing multiple genotypes, genotype was included as a covariate to ensure slope estimates were not confounded by differences among genotypes. A meta-analytic estimate of *α_plast:pℎyll_* was then calculated as the inverse-variance weighted mean of study-specific slope coefficients, weighting by the inverse of their squared standard errors.

For *L_em_*, we obtained a mean value of 6.5 from two studies in which *L_em_* was determined from plant dissections at emergence (Padilla & Otegui, 2005; Warrington & Kanemasu, 1983). This was consistent with reports showing four to five pre-formed leaves are present in the maize embryo (Bonnett, 1966; Kiesselbach, 1949; W.-Y. Liu et al., 2013; Padilla & Otegui, 2005), indicating that approximately two leaves are initiated between embryogenesis and emergence.

We compared seven different formalisms for predicting the number of visible leaves at T.I. (Table 1). These formalisms share a common general form but use different assumptions for model structure and parameterization. Model performance was evaluated based on standard metrics for prediction accuracy and error variance by comparing predictions with plant dissection data from our Night-Break platform experiment (described below). The following formalism was used for all subsequent analyses, with constants *α_plast:pℎyll_* and *L_EM_* derived from meta-analysis: *VL_TIi_* = *L_FLNi_* − *L_EM_*⁄*α_plast:pℎyll_* (Eq. 1); where, *VL_TIi_* is the visible leaf tip number at T.I. for the *i*^th^ gentoype, *L_FLNi_* is the final leaf number for the *i*^th^ genotype, *α_plast:pℎyll_* is fixed at 1.51 (dimensionless), and *L_EM_* is fixed at 6.5 leaves.

**Table 1.** Comparison of equations for predicting leaf tip number at tassel initiation.

| Study | Prediction Equation<sup>1</sup> | $R^2$ | RMSE<sup>2</sup> | MAE<sup>3</sup> | Bias<sup>4</sup> |
| --- | --- | --- | --- | --- | --- |
| Current study | $(L_{FLN_i} - 6.5)/1.51$ | 0.70 | 0.60 | 0.53 | -0.02 |
| Drouault et al., (2025) | $(L_{FLN_i} - 5.5)/1.73$ | 0.70 | 0.76 | 0.65 | 0.43 |
| Arnold (1969) | $0.46 \times L_{F_i}$ | 0.70 | 0.74 | 0.61 | 0.21 |
| Lejeune & Bernier (1996) | $(L_{FLN_i} - 1.95)/1.84$ | 0.70 | 1.22 | 1.05 | -1.04 |
| Tollenaar & Hunter (1983) | $0.50 \times L_{F_i}$ | 0.70 | 1.45 | 1.29 | -1.29 |
| Padilla & Otegui (2005) | $2.88 + [(L_{FLN_i} - 8.04) \times 0.63]$ | 0.70 | 1.65 | 1.54 | -1.54 |
| Arnold (1969) | $0.54 \times L_{FLN_i}$ | 0.70 | 2.13 | 2.04 | -2.04 |

<sup>1</sup>Some equations were algebraically rearranged from their original form to give visible leaf tip number as the common solution.
<sup>2</sup>Root mean square error: $\sqrt{[\sum(predicted - observed)^2/n]}$
<sup>3</sup>Mean absolute error: $\sum |predicted - observed|/n$
<sup>4</sup>Mean bias: $\sum(predicted - observed)/n$

### Experiments

This study combined data from a “Night-Break” experiment performed in 2024 in a greenhouse phenotyping platform, PhenoArch, hosted at the M3P (Montpellier Plant Phenotyping Platforms) (Cabrera-Bosquet et al., 2016), and two field experiments performed in 2023 in France and Mexico.

#### Platform experiment

Details for the Night-Break experiment were described previously (see Methods S2 in Drouault et al., 2025). Briefly, 10 genotypes (2369, BA90, B73, CML258, CML277, CML341, CML373, LH123Ht, Tzi8, and Tzi9) were tested under an 11/13-hour day/night cycle, without and with night-break (30s LED illumination per plant occurring between 21h00 and 23h00; beginning 13 days after sowing and applied each night for 21 days).

The Night-Break experiment was used to obtain ground truth data on the visible leaf tip number when T.I. occurs. To determine the timing of T.I. for each genotype-treatment combination, a total of 320 plants (16 plants of the 10 genotypes in each of the two photoperiod treatments) were examined through serial dissections of the shoot apical meristem (SAM), imaged using a Leica EZ4 dissecting microscope. T.I. timing was determined based on morphological changes of the SAM. First, a subjective scoring system based on the visual appearance of the SAM (Bonnett, 1966; Krüger, 1984) was used to classify the stage of its development. To define a more objective measure for estimating T.I. timing, we found that image-based quantification of the SAM’s length:width ratio (determined using ImageJ; Schneider et al., 2012) closely approximated the subjective scoring scale (**Fig S1**). Across genotypes and treatments, the visual score for T.I. timing corresponded to a mean SAM length-to-width ratio of 2.0, with a range among tested genotypes of 1.9 to 2.2 (**Fig S1**). Although a SAM length threshold has been used in some studies (Moncur, 1981; Vidal et al., 2021), a ratio is expected to mitigate length differences due to SAM size (Thompson et al., 2015). Therefore, the point at which the SAM reached an average length-width ration of 2.0 was used as the reference point for T.I. timing. Finally, the visible leaf tip number recorded on the same dissected plants were related to the SAM-based estimate of T.I., providing the visible leaf tip number (phyllochronic stage) at T.I. for each genotype-treatment combination (**Fig. S2**).

Additionally, for the two photoperiod treatments, eight random plants of each genotype were retained and grown to anthesis (total of 160 plants), then harvested to measure lengths of phytomer components and other traits (**Table S2**).

Although all measurements were recorded for all 10 genotypes, CML258 was excluded from the main study because of growth defects (buggy whipping) and late T.I., which introduced uncertainty into its T.I. estimate from serial dissection. Results for CML258 are shown in Supplementary Information. In addition, six plants were removed from the harvest data due to growth or measurement issues.

#### Field experiments

Two field experiments were carried out: one in Saint-Martin-de-Hinx, France (43.5704°, −1.3005°, planted June 8, 2023) and one in Puerto Vallarta, Mexico (20.81292°, −105.22242°, planted January 9, 2023). Both used a partial-replication design across two field sections (section 1 contained 160 plots with 114 genotypes, section 2 contained 320 plots with 256 genotypes) with independent randomizations of the same plot layout previously described for the Puerto Vallarta location (see Methods S2 in (Drouault et al., 2025). Plants were maintained under well-watered conditions throughout both trials.

The present study analyzed trait data from section 1 of each experiment, including blade lengths measured at multiple leaf ranks. Measurements were taken at the plant level, for up to four plants per plot. In each environment, blade lengths were measured at specific rank positions (the ear leaf, two ranks above the ear, and the penultimate leaf) in addition to a subset of leaf ranks spanning the canopy: 7, 8, 10, 14, 18, 20, and 23 in Mexico, and ranks 5, 9, 13, 17, and 22 in France (**Fig. S3**).

### Data Analysis

#### Comparing phytomer component lengths to tassel initiation timing

We used data from the platform experiment to compare the length profiles of each phytomer component (blade, sheath, and internode) in relation to timing of tassel initiation across individuals, which involved a multi-step process. First, individual plant profiles of blade, sheath, and internode lengths across rank positions were estimated by a generalized additive model with cubic regression splines and basis dimension k=11 (half the maximum final leaf number) using the mgcv package (Wood, 2011) in R (R Core Team, 2026). The function was extrapolated across a fine grid spanning the maximum range of leaf positions (from 1 to 22 in increments of 0.1), with predictions constrained to the final leaf number of each plant and truncated at zero to prevent biologically implausible negative values. Next, we converted the interpolated rank positions (blade, sheath, and internode numbers) into thermal times at which these structures formed. For leaf blades, we used the timing of leaf tip appearance, though we recognize this is an approximation since blades continue expanding after the tip first emerges. For sheaths and internodes, we used collar appearance because it better represents full elongation of these structures. We also converted the predicted visible leaf tip and collar numbers at T.I. to thermal time, allowing us to register the timing of interpolated phytomer profiles relative to T.I. across all plants.

#### Analysing phytomer profiles to compare canopy registration systems and decompose canopy variation

To address the alignment problem, we tested four registration systems for aligning blade-length data across plants: (i) absolute rank; (ii) position from T.I.; (iii) position from the dominant ear; and (iv) final leaf number normalization. Absolute rank aligns canopies by their absolute phytomer count (from 1 to *n*), while alignments by T.I. and dominant ear position are shifted according to phytomer ranks for each plant relative to those landmarks. Due to differences in final phytomer number, these registrations systems include flanking regions where plants differ in the common ranks that can be compared. In contrast, final leaf number normalization aligns canopies by their absolute count in proportion to their total number. This treats absolute rank differences in fractional terms while placing all plants on a common 0-1 scale. For each registration system, profiles were reconstructed for each plant using functional principal component (FPC) analysis via the PACE algorithm (Yao et al., 2005), implemented in fdapace (Zhou et al., 2024). Each plant’s trajectory was modeled as *X_i_*(*t*) = *μ*(*t*) + ∑*_k_ ξ_ik_φ_k_*(*t*) (Eq. 2); where, *μ*(*t*) is the mean profile, the *φ_k_* eigenfunctions estimated from the smoothed covariance surface, and *ξ_ik_* the corresponding scores. The mean function and covariance surface were estimated by kernel smoothing on a regular grid of 51 points, using the default proportional bandwidths (5% of the observation range for the mean and 10% for the covariance). FPC scores were estimated by conditional expectation, providing best linear unbiased predictions of each trajectory even for plants with few measurements (our data was from irregularly spaced leaves across the canopy, as mentioned above; **Fig. S3**). The number of components *K* was selected to explain 99.9% of the variance, and fitted profiles were evaluated across the canopy.

Modes-of-variation plots illustrating variance captured by each individual FPC were constructed synthetic profiles that deviate from the mean along a single eigenfunction (**Fig. S4**). For component *k*, each synthetic profiles were computed as 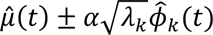 (Eq. 3); where, *α* is a scalar set to ±1 or ±2 standard deviations (corresponding to ±1 or ±2 standard deviations of the score 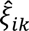) while remaining components were held at zero.

#### Estimating the influence of T.I. across functional principal components

To quantify the proportion of total functional variance attributable to T.I., a composite *R*^2^ was computed as the eigenvalue-weighted mean of the per-component *R* ^2^ values from regressing each FPCA score on T.I. This weighting ensures that components explaining more functional variance contribute proportionally more to the composite, giving a single interpretable measure of the linear association between T.I. and overall profile shape bounded between 0 and 1. A biplot of genotype-specific FPC1 and FPC2 scores illustrated the effect of T.I. on canopy shape variation, with arrows indicating the predicted trajectory across both components based on linear regression of T.I. on FPC scores for each field site.

#### Genome-wide association analysis

Genome-wide association analysis (GWAS) was performed using FarmCPU (X. Liu et al., 2016) implemented in the R package rMVP with the VanRaden genomic relationship matrix (Yin et al., 2021). This was performed separately for each field environment (France and Mexico). Genotype data was produced with the Maize 600k SNP array (Unterseer et al., 2014), which was reduced to approximately 370k SNPs in a merged dataset with a larger number of genotypes for other ongoing work (unpublished), and finally to approximately 300k SNPs after filtering for minor allele frequency at 10% within the set of test genotypes used for this study (genotype data was available for 110 of the 114 measured genotypes). Statistical significance was determined using a 1% Bonferroni-corrected *p*-value threshold to control for multiple testing.

#### Analysis of T.I.-independent scaling variation

Scaling variation independent of T.I., which we refer to as intrinsic scaling, was isolated by linear regression of the length of the longest leaf on T.I., using genotype mean values by environment (France, Mexico). Positive residuals indicated genotypes whose longest leaf was longer than expected given their T.I. (hyper-scalers), while negative residuals indicated genotypes whose longest leaf was shorter than expected (hypo-scalers). Genotypes were classified into three groups based on their residuals in each environment: consistent hyper-scalers (positive in both), consistent hypo-scalers (negative in both), and environment-specific scalers (opposite signs). Cross-environment agreement in intrinsic scaling among genotypes was determined by Pearson correlation between France and Mexico.

To estimate the genetic correlation of intrinsic scaling between environments, we fitted a bivariate mixed model treating France and Mexico as two correlated responses. The model included a fixed environment mean and a random genetic effect with an unstructured genetic covariance matrix **G**_0_ between environments, structured by the genomic relationship matrix **K**, and a diagonal residual covariance **R**_0_, allowing environment-specific residual variances. The genomic relationship matrix estimated as described above using a filtered set of approximately 113,000 SNPs, pruned by linkage disequilibrium with PLINK (v. 1.9.0-b.7.11; Chang et al., 2015). The model was fitted by REML in sommer (v. 4.4.5; Covarrubias-Pazaran, 2016). The genetic correlation was obtained as: 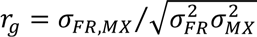, and the line-mean heritability within each environment as 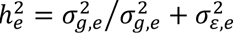

## Results

### A conserved plastochron–phyllochron relationship estimated across studies improves prediction of T.I. timing

The plastochron–phyllochron ratio, *α_plast:pℎyll_*, a measure of the relative timing of leaf primordia initiation versus leaf tip appearance, is fundamental to equations predicting maize development, including the timing of tassel initiation (T.I.) examined here. We estimated this ratio by meta-analysis of 11 studies, using the regression of leaf primordia number on leaf-tip number to obtain the slope of the relationship per study. Studies varied substantially in total observations (*n*=15 to 590). To account for this, we computed the inverse-variance weighted mean of the study-specific slopes, yielding a *α_plast:pℎyll_* of 1.51 ± 0.01 (Figure 2A); study weights ranged from 1% (Tollenaar & Hunter, 1983, with 16 observations available from Padilla & Otegui, 2005) to 66% (Padilla & Otegui, 2005, with 590 observations). Because Padilla & Otegui (2005) included 19 genotypes (all other studies had no more than two), we used it to test for genotypic variation in *α_plast:pℎyll_*. Within that study, genotype had no significant effect (*p* = 0.54), whereas *α_plast:pℎyll_* differed significantly across studies (*p* < 0.001). Indeed, the range in *α_plast:pℎyll_* across studies (1.13 to 1.86) was greater than the range among genotypes (1.38 to 1.62).

**Figure 2.**
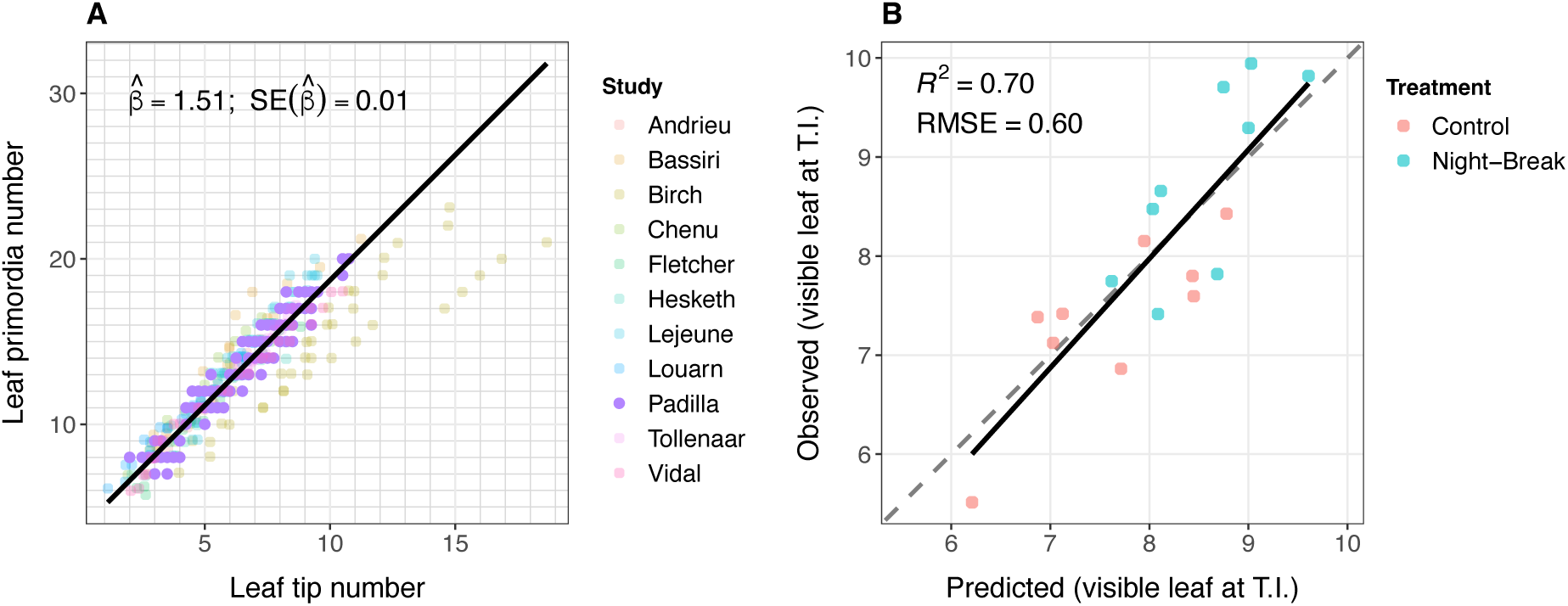
Meta-analytical parameter estimation for predicting leaf tip number at tassel initiation in maize. **(A)** Relationship between the number of leaf initiation primordia and visible leaf tips across studies to derive *α_plast:pℎyll_*. Data from each study is shown as separate colors with the first author of the study indicated in the legend. Color intensity reflects the inverse error variance weighted contribution of each study toward the pooled slope estimate (e.g., the Tollenaar study had 1% weight while the Padilla study had 66% weight). The pooled slope 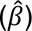 and standard error (SE) estimate for *α_plast:pℎyll_* is indicated and the fitted line shown (solid line). **(B)** Relationship between visible leaf tip number at tassel initiation for observed (determined by serial plant dissections) and predicted (Eq. 1) values, based on data for 10 genotypes in the Night-Break platform experiment, with photoperiod treatments shown as separate colors. The adjusted *R*^2^ and root mean square error (RMSE) from a linear regression are indicated, as are the fitted (solid) and 1:1 (dashed) lines.

To evaluate our meta-analysis results, we compared observed and predicted values of 10 genotypes measured under two photoperiod treatments in the Night-Break platform experiment, with constants for *α_plast:pℎyll_* = 1.51 and *L_em_* = 6.5 leaves (both from the meta-analysis). Predictions agreed well with meristem dissection-based estimates of T.I. (*R*^2^ = 0.70; **Fig. 2B**), which further supported the conservation of *α_plast:pℎyll_* across genotypes. **Table 1** summarizes prediction accuracies and error variation for Eq. 1 compared to several other equations that use different constants or formalisms. All equations had the same predictability but varied by their degree of error and bias. The common formalism from Lejeune & Bernier (1996) and Drouault et al. (2025) substituted with constants obtained through our meta-analysis performed best overall (**Fig. 2B**, **Table1**). Therefore, throughout this study we used this to predict T.I. timing.

### Timing of floral transition structures length profiles of the maize canopy

Prior research has proposed that at the individual plant level, the maize canopy is formed through coordinated development of phytomer components, with distinct size scaling effects during the vegetative (before T.I.) and reproductive (after T.I.) phases (Zhu et al., 2014). If this coordination is conserved across genotypes and environments, variation in canopy size would arise systematically from variation in T.I. timing. Under this model, we expect phytomer size to scale across phytomer ranks with the timing of T.I. (**Fig. 1**), and the rank position of the dominant ear (uppermost axillary bud) to scale in parallel. To test this, we leveraged platform and field experiments to compare profiles of phytomer lengths among plants where the predicted phytomer rank formed at T.I. varied. Across genotypes and photoperiod treatments (platform) or environments (field), the number of phytomers formed at T.I. ranged by five in the platform experiment and 12 in the field experiment.

First, we used blade, sheath, and internode length measurements from the platform experiment (**Table S2**) to examine how each component’s length changed with the timing of tassel initiation. For each plant, we linked the length measurements at each rank position to the thermal time when that rank was formed, allowing us to relate the developmental timing of individual phytomers in relation to T.I. timing (using the corresponding leaf tip rank for blade size comparison and the collar rank for sheath and internode size comparison; see Methods). The results revealed that while each phytomer component displayed a distinct profile shape, these patterns remained broadly consistent across different genotype and photoperiod treatments (**Fig. 2A**). Overlying the overall mean profiles of phytomer component lengths relative to T.I. timing, we captured their coordinated developmental sequence: blade and sheath components of the leaf elongated nearly simultaneously (blades elongated slightly sooner) and corresponding internodes elongated afterward (**Fig. 2B**). The leaf components and internodes reached their peak lengths at progressively later times from T.I., before each was reduced in size.

Using data from both platform and field experiments, we also tested the expectation that T.I. is linked to the rank of the dominant ear. Predicted values for T.I. explained 69% and 90% of the variation in observed dominant ear rank for the platform and field experiments, respectively. Ear rank, often used as a reference point in maize phenotyping, correlated with cardinal points of the phytomer components similarly to T.I. (**Table 2**, platform data), indicating that it provides a close approximation of tassel initiation. We return to this relationship in the next section.

**Table 2.** Correlations between tassel initiation and dominant ear position with cardinal points of phytomer component profiles.

| Reference developmental point<sup>1</sup> | Max blade rank | Max sheath rank | Sheath plateau rank<sup>2</sup> | First internode rank<sup>3</sup> | Max internode rank |
| --- | --- | --- | --- | --- | --- |
| T.I. | 0.71*** | 0.36*** | 0.75*** | 0.23** | 0.20* |
| Ear position | 0.76*** | 0.49*** | 0.76*** | 0.26** | 0.11<sup>ns</sup> |

<sup>1</sup>Results from the platform experiment. Pearson correlations coefficients significant at \*\*\*p < 0.001, \*\*p < 0.01, \*p < 0.05, ns= not significant
<sup>2</sup>Sheath plateau rank identified as the changepoint (Killick & Eckley, 2014) marking the shift from declining to stabilized inter-leaf size, or the rank of minimum absolute decline if no changepoint was detected.
<sup>3</sup>First internode rank corresponds to the first elongated internode that could be measured (not the true first internode).

### Leaf-number normalization provides the most effective registration for canopy comparison

Having used T.I.-aligned canopies from the platform experiment to describe the structure of phytomer-component profiles, we next asked whether T.I. alignment is the best approach for comparing canopy variation. For this, data from the field experiments (France and Mexico) were used, where broader variation in final leaf number and T.I. provides a more discriminating test. We compared four methods of phytomer registration: (i) absolute rank; (ii) position from T.I.; (iii) position from the dominant ear; and (iv) final leaf number normalization. These experiments measured only blade length (not sheaths or internodes) and only across a subset of phytomer ranks (**Fig. S3**). We therefore applied functional principal component analysis via PACE (Yao et al., 2005) to reconstruct full blade-length profiles. We compared registration methods by examining the overall mean and the correlation structure of blade-length profiles across registration coordinates, and quantifying residuals between observed and predicted points in the reconstructed profiles.

Registration systems influenced the mean blade-length profile and structure of covariation across the grid of interpolated coordinates for phytomer ranks, as shown in **Fig. 4A-B** with the windows for T.I. timing highlighted as grey boxes. With absolute rank alignment, the wide window for T.I. (spanning 4-16 leaves across genotypes and environments) reflected how blade sizes at each rank position corresponded to leaves formed at different times relative to tassel initiation. The T.I.-window size was reduced in alignments based on the dominant ear rank and normalized leaf number, with mean profiles that were more similar to T.I.-based alignment, and a peak blade length position congruent with complete measurements in the platform experiment (right-flanking T.I.; cf. **Fig. 3A**). Compared to absolute rank, the three phenological registration systems also produced pairwise correlations in blade lengths across coordinates that were more positive overall with higher correlation persisting at greater lag distances (**Fig. 4B**), reflecting the conservation in profile shape across plants. Based on the analysis of residuals, however, leaf number normalization yielded substantially better blade-length profile reconstructions than the other registration systems, with an overall reduction in RMSE by 40–50% and residuals that were consistently unbiased across the registration coordinates (**Fig. 4C**).

**Figure 3.**
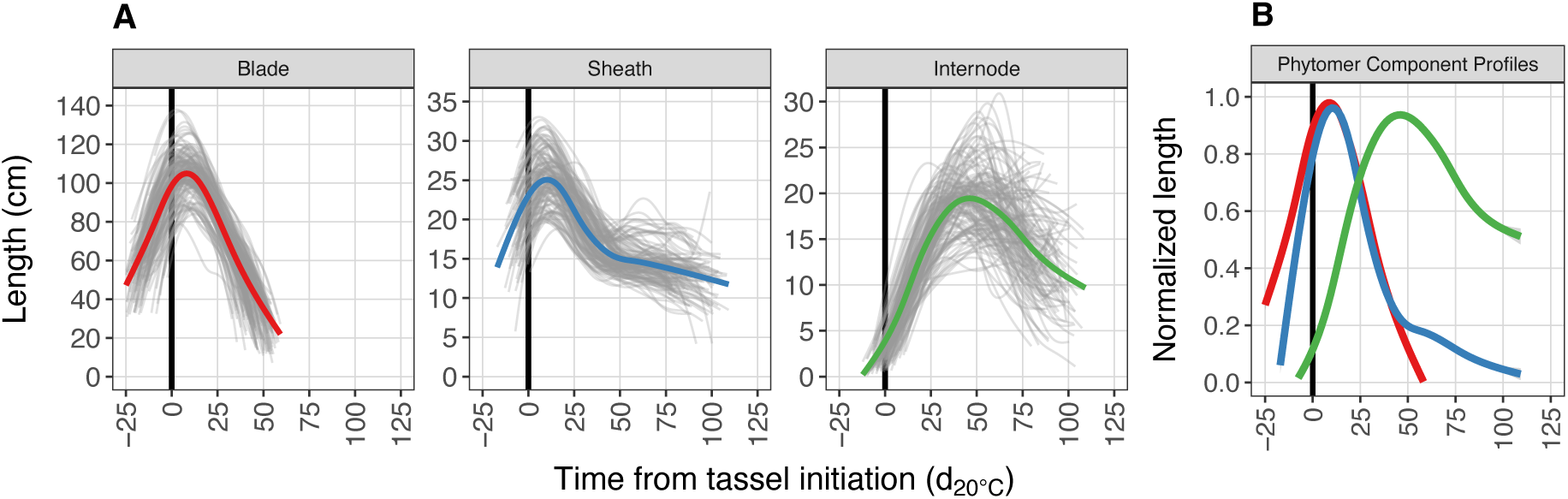
Length profiles of phytomer components in relation to thermal time from tassel initiation. **(A)** Phytomer component length profiles by plant (grey lines; *n* = 142, 9 genotypes) from the platform experiment. For each component (blade, sheath, and internode), the length measured at each rank position along the shoot was related to the formation of the corresponding rank in thermal time relative to tassel initiation (T.I., marked by the black vertical line). Rank-to-time registrations were based on leaf tip number for blades and collar number for sheaths and internodes. The overall mean profile is shown for the blade (red), sheath (blue), and internode (green). **(B)** Length normalized profiles from (**A)** combined in a single plot, with phytomer components labeled by their corresponding color.

**Figure 4.**
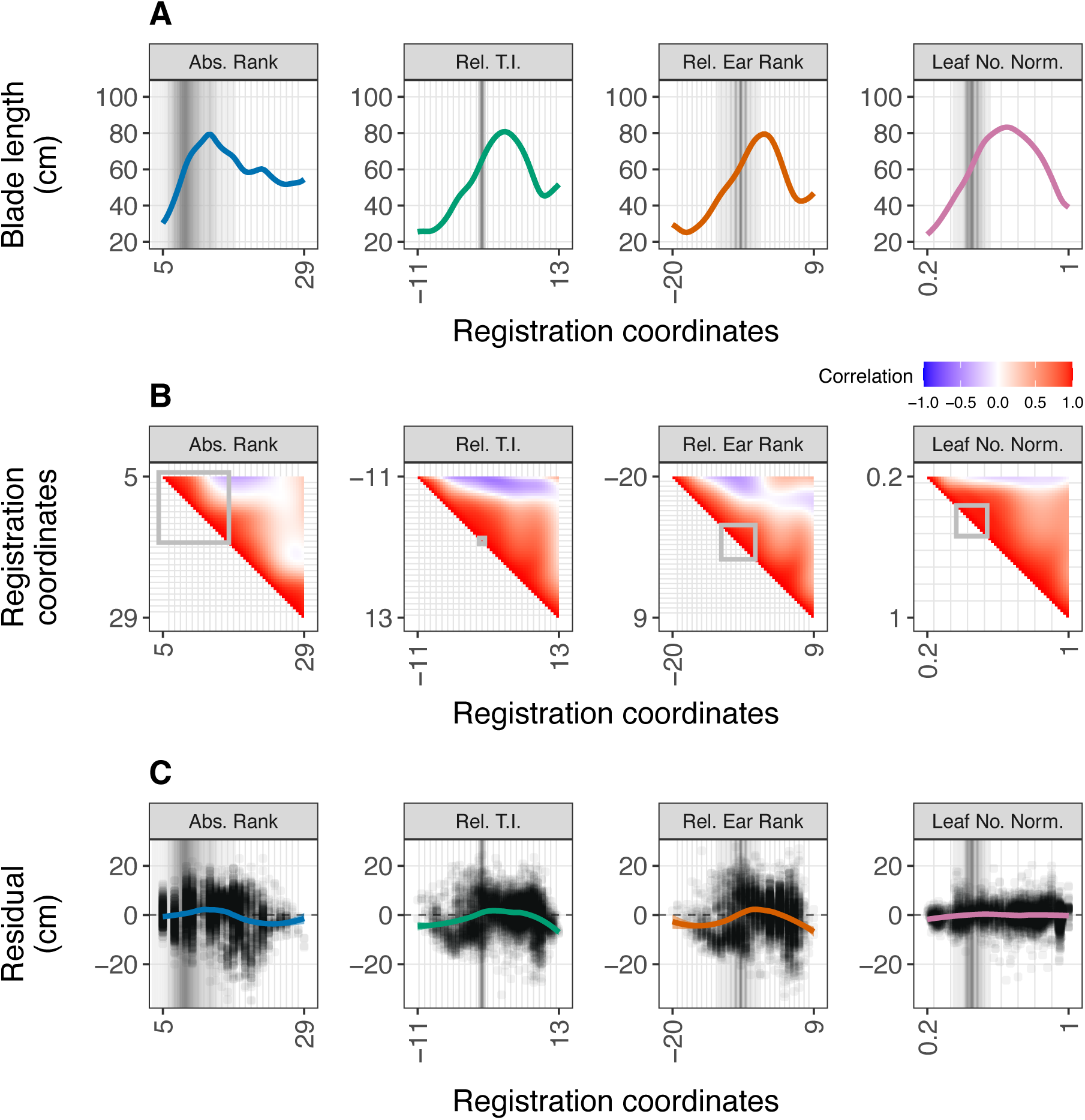
Effect of registration systems on size variation in leaf blade-length profiles. Four registration systems (facet labels) were used to align phytomer ranks for pooled data from both field experiments (*n* = 879 plants across environment and genotype treatments). (**A**) Mean blade-length profile size curves across coordinates for each registration system (x-axes). (**B**) Pairwise correlations in blade-length size between registration coordinates. (**C**) Residual deviations in predicted blade-lengths (cm) across registration coordinates. Each point represents the residual deviation for an individual leaf length measurement from the corresponding plant-specific prediction; the smoothed line and shaded band show the loess trend with 95% confidence interval. Grey shaded regions indicate the density of predicted tassel initiation (T.I.) across all plants for each registration system.

Together, these results showed that aligning by absolute rank inadvertently compares phytomers formed at different developmental stages across plants, conflating size variation with phase differences in T.I. timing. The three phenological registration systems help to control for this, with each exposing the canonical shape of the maize blade-length profile, where blade sizes increase until T.I. and decrease approximately thereafter (cf. **Fig. 3A** and **Fig. 4A**). While registration relative to T.I. timing or ear rank were successful in recovering the conserved shape of blade-length profiles, these alignments retain phytomers that are incomparable between plants (due to the differences in absolute number). This leads to biased reconstruction in flanking regions of the blade-length profile, which tends to increase with distance from the reference point. However, for blade-length profile reconstruction, normalizing by final leaf number densifies data coverage across registration coordinates, yielding the most accurate PACE-based reconstructions with the lowest error and no apparent rank-dependent bias.

### A persistent effect of floral transition timing partially explains scaling variation in canopy size

With leaf-number normalization established as the most effective registration system for comparing blade-length profiles, we used PACE to: (i) characterize how these profiles vary in shape and (ii) how much of that variation is explained by tassel initiation.

Functional principal component (FPC) analysis of profiles from the field experiment, with all plants from both environments analyzed together, showed that variation in profile shape arose from two main components. Amplitude variation reflects the overall size of the profile, and phase variation reflects horizontal shifts in the position of features along it. The first two FPCs captured over 90% of this variation. Modes-of-variation plots (**Fig. S4**; see Methods) showed that FPC1 (77%) was dominated by amplitude, a vertical shift above and below the mean curve reflecting scaling in blade length, with a smaller phase component of about one leaf-normalized unit. FPC2 (16%) was dominated by phase, with overlapping curves shifted horizontally by about two leaf-normalized units, reflecting variation in the position of the longest blade. The consolidation onto a single dominant axis was apparent across all three phenological registration systems (**Fig. S4**). Under absolute-rank registration, by contrast, variance was distributed more evenly between FPC1 and FPC2 (50% and 35%, respectively), and the modes of variation appeared highly conflated, confirming that good registration is necessary not only for accurate reconstruction but also for interpretable decomposition.

To examine these interpretations quantitatively, we extracted two features from individual profiles: the longest blade length as a scaling proxy, and its rank position as a phasing proxy. Because reconstructed curves integrate contributions from all FPCs, correlations between these proxies and individual FPC scores reveal which FPC is the dominant driver of each feature. The scaling proxy correlated strongly with FPC1 (Pearson *r* = 0.97) and was uncorrelated with FPC2 (*r* = 0.02), cleanly isolating the scaling mode of variation. The phasing proxy correlated strongly with FPC2 (*r* = 0.85) and moderately with FPC1 (*r* = −0.49). Unlike the scaling proxy, which was uncorrelated with FPC2, the phasing proxy retained a partial correlation with FPC1, consistent with FPC1’s smaller phase component indicated above. Together, the modes-of-variation analysis and the proxy correlations provide a coherent decomposition of canopy shape variation: a dominant scaling mode coupled with a smaller phase shift (FPC1), and a secondary phase-dominant mode (FPC2).

We next asked how much of this shape variation was associated with T.I. timing. Two complementary analyses converged on the same answer. First, total variation across all FPCs calculated by a composite *R*² statistic showed that T.I. explained 35% of the variation in blade-length profiles for the pooled data (split by environment, composite *R*² = 28% in France and 8% in Mexico), which was dominated by FPC1, with negligible contributions from subsequent components (**Fig. 5A**). This was reflected in a biplot of FPC1 and FPC2 (**Fig. 5B**), where the vector representing T.I. timing aligned with the FPC1 axis in both field environments, confirming that T.I. variation acts primarily as a scaling effect on blade-length profiles with a stronger effect in France. Second, in the long-day environment of France (but not in the short-day environment of Mexico) where the T.I. association was stronger, GWAS of FPC1 scores and predicted T.I. timing identified the same genetic association with *ZmCCT10* (**Fig. 5C**), a central gene for T.I. timing in maize affected by photoperiod conditions (Hung et al., 2012; Yang et al., 2013). The environment-specific signal is consistent with *ZmCCT10*’s known role in suppressing T.I. under long days, which leads to the formation of additional leaves and in turn a larger scaling of blade-length profiles. This provided independent genetic support for a functional link between T.I. timing and blade-length scaling. Together, these analyses indicate that T.I. timing acts as a persistent driver of blade-length scaling variation. This is consistent with a developmental scaling effect in which forming more leaves before the reproductive transition yields a larger maximum blade, shaping canopy size differences even where scaling explains only part of the total shape variation observed.

**Figure 5.**
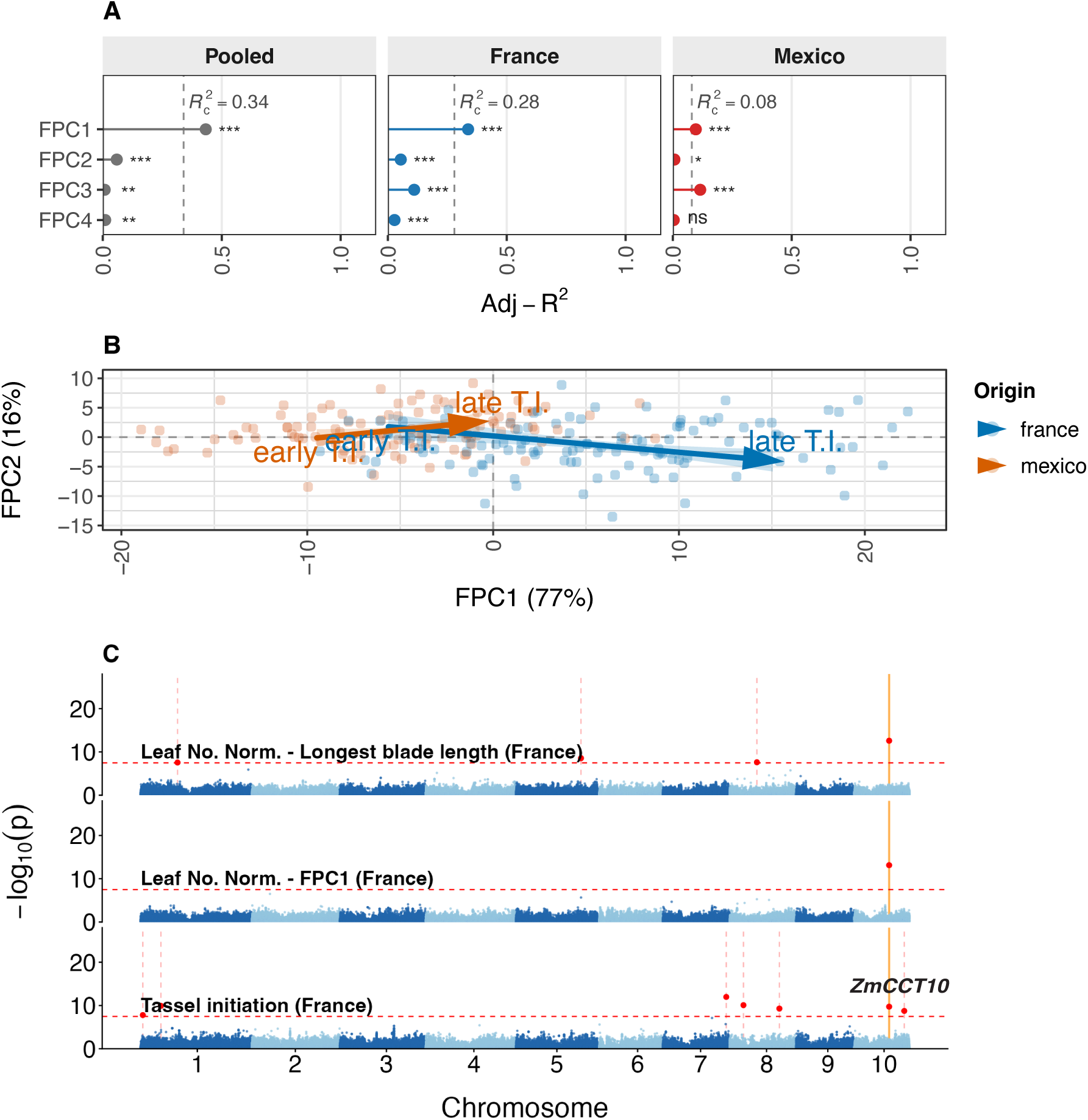
Associations between blade-length profile variation and tassel initiation. (**A**) Amount of variation in individual functional principal components (adjusted *R*^2^ per FPC) and their total variation (*R*^2^) explained by tassel initiation. The faceted plots show results from the pooled analysis and for each of the two field locations. (**B**) Biplot of scores for the first two FPCs from analysis of the pooled data with the average predicted trajectory of T.I. indicated per environment. The functional variation in blade-profiles attributed to each FPC is indicated in parentheses. Plot aspect ratio was scaled to reflect the proportion of variance explained by FPC1 relative to FPC2. (**C**) Manhattan plots of GWAS results for T.I. timing (tassel initiation), FPC1, and length of the longest leaf blade from the field experiment in France. The dashed horizontal line in red indicates the Bonferroni-corrected threshold at alpha 0.01. Red points marked with dashed vertical lines mark significant associations, and orange solid lines mark common associations across traits. The one association common to all traits on chromosome 10 is present within the *ZmCCT10* gene (Zm00001eb418700, B73 NAM reference position 96,095,110).

### Intrinsic scaling represents a separate axis of variation for manipulating canopy size

The partial relationship between T.I. timing and longest blade length suggests that not all scaling variation is explained by the T.I.-linked effect of final leaf number. The remaining variation, which we term intrinsic scaling, may underlie plant productivity and serve as a target for selection in breeding. We quantified this intrinsic component by taking the residuals of longest blade length after regressing on T.I. timing within each environment; FPC1 gave equivalent results, consistent with its strong loading on blade length.

Correcting longest blade length for T.I. timing removed 34% of its variance in France and 16% in Mexico, leaving substantial residual variation in both environments. This residual was positively correlated across environments (*r* = 0.67), indicating that a genotype’s tendency to over- or under-scale relative to its T.I. timing is partly conserved across contrasting environments (**Fig. 6**). Forty-two genotypes scaled consistently above expectation in both environments (hyper-scalers) and 42 consistently below (hypo-scalers), while 24 showed environment-specific scaling. Estimating the cross-environment genetic covariance of intrinsic scaling indicated a heritable trait (ℎ^2^ = 0.85 and 0.92 in France and Mexico) with a strong genetic correlation between environments (*r*_g_ = 0.79). Together, these results identify a heritable-candidate axis of canopy scaling that is decoupled from phenology and may be accessible to selection.

**Figure 6.**
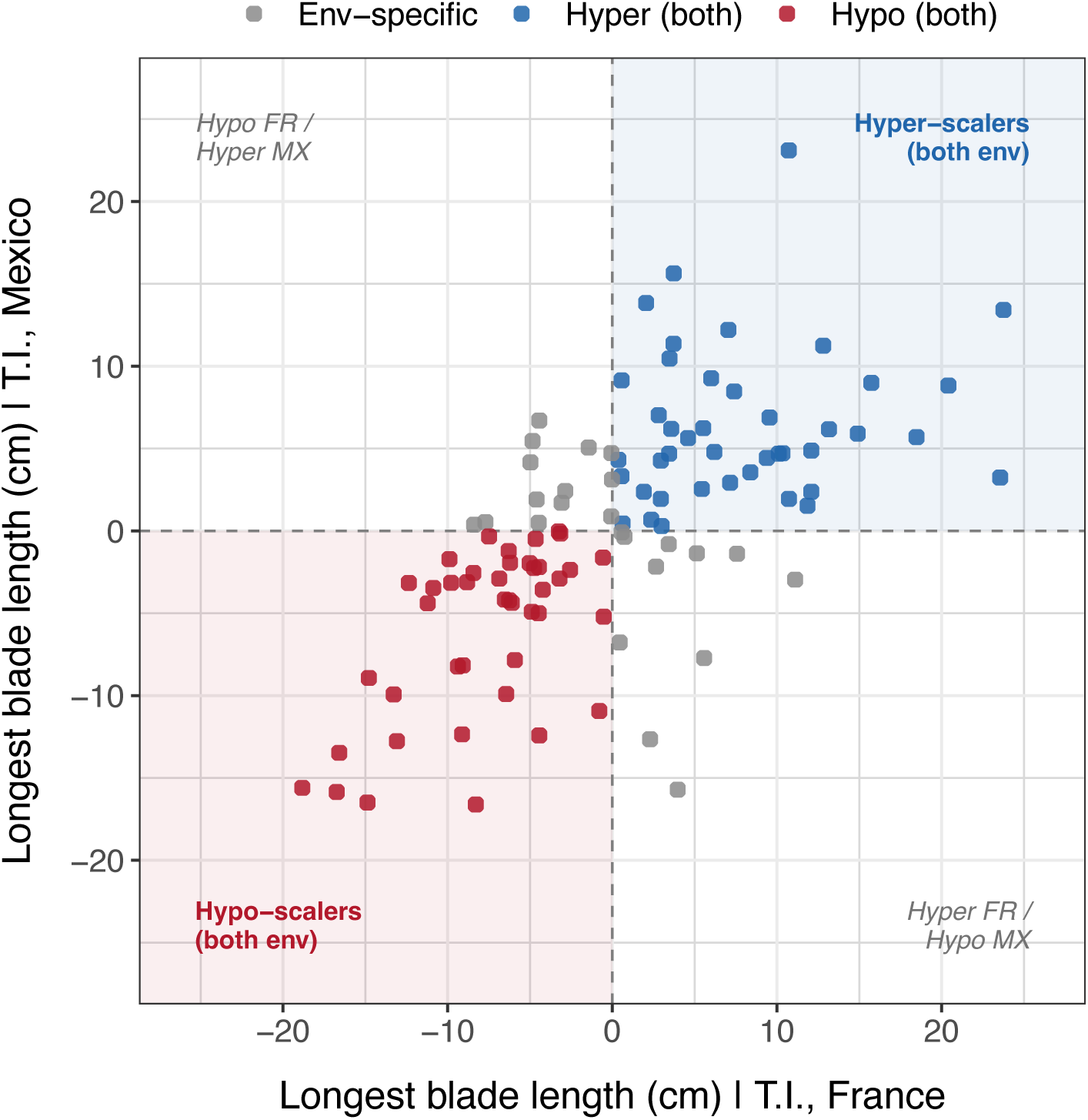
Intrinsic scaling variation in longest blade length. Each point shows a genotype’s mean residual deviation in longest blade length (cm), computed as the residual of longest blade length regressed on T.I. timing within each environment. Axes give the residual in France (x) and Mexico (y). Shaded quadrants mark genotypes that scaled consistently below (hypo-scalers, lower-left) or above (hyper-scalers, upper-right) the value expected from their T.I. timing in both environments; off-diagonal quadrants contain genotypes with environment-specific (contrasting) scaling. Residuals were correlated across environments (*r* = 0.67; *n* = 108 genotypes).

## Discussion

In comparative biology, a recurring challenge is how to make meaningful comparisons across individuals that differ in size and developmental timing (Barton, 2024). This is becoming more important as high-throughput phenotyping increasingly captures temporal and profile data, where each individual is measured as a developmental trajectory rather than a single value (Miao et al., 2020; Mu et al., 2022; Sweet et al., 2024; Xu et al., 2018). Comparing canopy architecture poses exactly this problem: plants differ in total phytomer number, and phytomer size scales with that number, so a naive comparison by absolute rank conflates distinct sources of variation (**Fig. 1**). Ultimately, we show that expressing phytomer-component profiles relative to final phytomer number resolves this for canopy comparison, but to reach that conclusion we first developed an improved, broadly calibrated predictor for T.I. timing (**Table 1**, **Fig. 2**), then examined how dimensioning of the shoot system is coordinated around the reproductive phase change marked by tassel initiation (**Fig. 3**).

In maize, the effect of developmental kinetics and final leaf number on phytomer-component size has been studied for the blade (Lacube et al., 2020; Vidal et al., 2021), sheath (Andrieu et al., 2006; Vidal et al., 2021), and internode (Birch et al., 2002), but T.I. timing itself has not previously been positioned as a landmark within the developmental timeline of the vegetative shoot system. By mapping the length profiles of all three components against the thermal time of their formation, relative to T.I., we show how the shapes of these profiles anchor to a single meristem event (**Fig. 3**). Because T.I. occurs while several phytomers are still being formed or are actively developing, this reveals where each component’s dimensioning shifts relative to the transition, as well as the temporal coordination in scaling among components. For the leaf blade and sheath, the profile peaks fell shortly after T.I. (approximately 8 days at 20 °C), with subsequent leaves progressively downscaling to give a characteristic bell shape (**Fig. 3**). In contrast, internode profiles showed elongation starting at tassel initiation. Based on this temporal proximity, we hypothesize that T.I. triggers these changes, acting through an acceleration in blade–sheath boundary formation described previously (Birch et al., 2007; Vidal et al., 2021). The transition to T.I. and an acceleration in boundary formation may form a single causal sequence, with T.I. as the upstream signal regulating bud outgrowth as proposed by McSteen & Leyser (2005), which combines with a shorter path length for leaf tip appearance due to rapidly extending internodes as suggested by Skinner & Nelson (1995). The ear position along the stem was similarly correlated with cardinal points in the phytomer profiles (**Table 2**), consistent with synchronized changes in the shoot apical meristem (T.I.) and axillary bud commitment to ear formation. It is unclear whether the centrality of T.I. in shoot organization is specific to maize due to its two axes of reproductive development (tassel and ear), compared to most other grasses that form a sole panicle (Schrager-Lavelle et al., 2017).

As T.I. in maize is a single developmental event that organizes canopy shape and is shared across plants, it provides a natural reference point for comparison. Using functional data analysis (Yao et al., 2005), we therefore compared phenological registration systems for aligning blade-length profiles (anchoring to T.I., to ear rank, or normalizing by final leaf number) against naive alignment by absolute rank (**Fig. 4**). The phenology-based systems better reflected the expected profile structure (**Fig. 4A, B**), but only normalizing by final leaf number reconstructed profiles without bias (**Fig. 4C**). Residuals were nearly centred on zero across the full normalized scale (minor deviation for the lower canopy), whereas anchoring to T.I. or ear rank left systematic prediction bias across ranks.

This bias is not merely a loss of accuracy. Misalignment lets phase variation (where features fall along the profile) contaminate amplitude variation (overall size), a well-recognized problem in functional data analysis (Marron et al., 2015). Anchoring to T.I. or ear position illustrated this cost. Shifting profiles to a common landmark improved local alignment but created new mismatches and gaps at the extremes, where varying numbers of phytomers fell on either side of the landmark. This distorted the reconstructed profiles, evidenced by bias in the fit (**Fig. 4C**). Because this distortion was conditioned by the registration landmark itself, it was absorbed into the FPC scores as apparent T.I.-related variation, inflating their association and mirroring the confounding seen under absolute-rank registration (**Fig. S5**). By reconstructing without positional bias and concentrating variation in the fewest components, leaf-number normalization separated shape and size variation more cleanly. This helps to resolve the alignment problem and provides a generalizable solution for comparing canopy architecture across diverse genotypes and environments.

In addition to addressing the alignment problem, we reveal an ontogenetic link between T.I. timing and scaling variation in maize blade-length profiles. T.I. plays a dual role. It organizes the conserved shape of the canopy profile while also scaling its size with leaf number, contributing to variation in blade length across genotypes and environments. In a long-day environment, photoperiod suppresses the floral transition to differing degrees across genotypes, from genotypes that are insensitive to those that are strongly sensitive, widening the variation in final leaf number (Bonhomme et al., 1991; Drouault et al., 2025; Ellis et al., 1992). Because genotypes diverge in how far T.I. is delayed, and blade length scales with the number of phytomers formed at T.I., the genotypic variation in T.I. timing becomes a driver of blade-length scaling. We found that this accounted for roughly a third of the scaling variation in a long-day environment of France. The genetic association with maize *ZmCCT10 (Ghd7)*, a central gene in the photoperiod-sensing network for flowering time in cereal crops (Murphy et al., 2014; Stephenson et al., 2019; Xue et al., 2008), reinforced this link and was specific to this environment. Its detectability in our modest panel (*n* = 110) is consistent with the larger effect of this locus on maize flowering-time variation in long-day environments (Hung et al., 2012; Stephenson et al., 2019; Yang et al., 2013). By contrast, in a short-day environment (Mexico) the T.I.-linked effect explained little scaling variation and no genetic associations were detected (**Fig. 5**).

Decoupling the phenological effect of T.I. isolated a separate source of intrinsic-scaling variation in blade length, independent of T.I., as a candidate for genetic dissection and breeding. This residual axis has a genetic basis evidenced by high heritabilities (ℎ^2^ > 0.85) in both environments and a strong genetic correlation between them (*r*_g_ = 0.79), enabling consistent hyper- and hypo-scaling genotypes to be distinguished (**Fig. 6**). However, we detected no associations with intrinsic scaling in this study, which we attribute to its likely polygenic architecture, beyond the mapping power of our modest panel. Nonetheless, our results indicate a heritable component of scaling variation that warrants dissection in larger populations. This variation could be used to drive selection for smaller or larger canopies while holding T.I. timing constant within a specific environmental context, decoupling canopy size from phenology. A further axis, related to longest-leaf position (the phase variation captured by FPC2), could be selected to redistribute the longest leaf within the canopy, reshaping the profile independently of its size. This opens new opportunities for manipulating canopy architecture for crop improvement.

### Conclusions

This study advances our ability to compare maize canopy architecture by grounding it in development, revealing how the canopy is organized around the floral transition. When this is overlooked, sources of variation become conflated and may lead to misinterpretation. The developmentally adjusted, functional data analysis demonstrated here for whole-plant blade length variation could be extended to other leaf traits, such as width or erectness (Elli et al., 2023), or applied equally to sheath and internode profiles. Co-localizing these component profiles on a common developmental time axis across individuals (**Fig. 3**) offers a new way to study conservation and deviation in the patterned growth of maize. Developing this approach for other grasses (Poaceae) could lead to a general framework for comparing canopy architecture within and between species.

By disentangling shape and size variation in blade-length profiles, we defined a distinct axis of canopy variation independent of the timing of floral transition. Our findings indicate that this represents a sizeable fraction of size variation (65% or more), including a genetic component that drives consistent hyper- and hypo-scaling of blade length across environments (**Fig. 6**). This is relevant to crop improvement as a selection target for modifying canopy architecture, decoupled from phenology. It may reflect differences cell division and expansion rate (Baute et al., 2015; Gonzalez et al., 2010; Tardieu et al., 1999; Vidal et al., 2021), though dissecting the specific mechanisms and the genetic basis requires further work.

Finally, the phenological registration required for meaningful comparison of shape and size exposes a deeper problem. Even when phytomers from different individuals in the same environment are matched by developmental stage, that stage is reached at different times across individuals, so the phytomers develop under different environmental conditions. Comparisons of profile size, or of size at a given position, are therefore further confounded with these temporal differences in environment. This represents an ontogenetic-by-environment interaction that is rarely recognized in plant science, yet it limits how cleanly genetic and environmental contributions can be separated. Recognizing this interaction is a step toward isolating the more intrinsic sources of genetic variation in plants.

## Acknowledgements

Experiments on the HTP-platform in Montpellier, France, were made possible thanks to the M3P team, including Llorenç Cabrera-Bosquet, Maëlle Lis, Aurélien Besnier, Nora Picaut, Noé Lalouette-Marrier d’Unienville, Anthony Rosello, Javier Soto Mendoza, and Aurélien Ausset. Field trials were conducted with the support of Arnaud Brunet, Bernard Lagardère, and Jean-René Loustalot from the experimental station at Saint-Martin-de-Hinx, France, as well as the Moreno Retis team, including José Moreno and Luis Rodríguez, from trials in Puerto Vallarta, Mexico. We thank María Otegui and Juan Padilla from the University of Buenos Aires for sharing plastochron and phyllochron data used for the meta-analysis. We also thank Delphine Madur and Valerie Combes at INRAE (GQE–Le Moulon) for their collaboration on the production of genotype data.

## Declaration of AI-assisted technologies in the writing process

During the preparation of this manuscript, Claude (Anthropic) was used to assist with refining the wording and clarity of the text, and for developing R code for data analysis and visualization, which the authors executed and verified.

## Data Availability

Datasets and computer code used for this study are publicly available in the cancom repository hosted on the Maize ATLAS GitHub site (https://github.com/maizeatlas/cancom)

## Author Contributions

KJ, CF, RJW: conceptualization; KJ, CF, CP investigation; KJ, CF, RJW: methodology; KJ, RJW: formal analysis; KJ, RJW: writing; RJW: funding acquisition.

