## Supplementary Information for "Disentangling shape and size variation to compare canopy architecture in maize"

The following supporting information is available for this article:

**Method S1.** Function used for thermal time estimation.

**Table S1.** Summary of studies on plastochron and phyllochron rates in maize.

**Table S2.** Traits measured in platform and field experiments.

**Figure S1.** Comparison between methods for determining tassel initiation of the shoot apical meristem.

**Figure S2.** Relationship between the visible leaf tip and development of the shoot apical meristem.

**Figure S3.** Observation density for leaf blade length measurements from the field experiment.

**Figure S4.** Modes-of-variation for functional principal components by registration system.

**Figure S5.** Variation in functional principal components explained by tassel initiation.

### Method S1. Function used for thermal time estimation.

Thermal time was used in the estimation of phyllochron (leaf tip appearance) and collochron (collar appearance), and the timing of tassel initiation. This was calculated using the following equation from Parent & Tardieu (2012) :

$$f(A, T_h, \Delta H_A{}^{\ddagger}, R, \alpha, T_0) = \frac{A T_h e^{\left(\frac{-\Delta H_A{}^{\ddagger}}{R T_h}\right)}}{1 + \left[e^{\left(\frac{-\Delta H_A{}^{\ddagger}}{R T_h}\right)}\right]^{\alpha(1-\frac{T_h}{T_0})}}$$

$A$ is a scaling coefficient fixed at 5.16 x $10^{10}$, $T_h$ is hourly temperature (°K), $\Delta H_A{}^{\ddagger}$ (J $mol^{-1}$) and $T_0$ (°K) are shape parameters specific to each species (73900 J $mol^{-1}$ and 306.4 °K), $R$ is the universal gas constant (8.314 J $K^{-1}$), and $\alpha$ is a constant valid for most species fixed at 3.5 (dimensionless).

Parent, B., & Tardieu, F. (2012). Temperature responses of developmental processes have not been affected by breeding in different ecological areas for 17 crop species. *New Phytologist*, 194(3), 760–774. https://doi.org/10.1111/j.1469-8137.2012.04086.x

### Table S1. Summary of studies on plastochron and phyllochron rates in maize.

| Study[1] | Genotypes | Experimental conditions | Definition of leaf initiation | Instrument | Data location |
|---|---|---|---|---|---|
| Thiagarajah & Hunt (1982) | A498 x CG10 (Hybrid) | Growth chamber; Contrasting day/night temperatures: 30/25°C, 25/20 °C; 15h | Visible to the naked eye | None | Fig.1, pg.1648 |
| Tollenaar & Hunter (1983) *, extracted from Padilla & Otegui (2005) | Guelph GX122 (Hybrid) | Growth chamber; Photoperiod treatments: 10h vs. 20h | Not stated | Binocular microscope | Fig.4(A), pg.1004 |
| Warrington & Kanemasu (1983) | DeKalbXL45 (Hybrid) | Growth chamber; 26/21°C;12h | A ridge visible flanking the apex | Microscope | Fig.1 and Fig.2, pg. 757 |
| Hesketh & Warrington (1989)* | DeKalbXL45 (Hybrid) | Growth chamber; 26/21°C;12h | Length of the apex ridge | Stereomicroscope | Fig.2, pg.697 |
| Zur et al. (1989) | P3377 (Hybrid) | Growth chamber; Contrasting day/night temperatures: 26/20°C, 30 /24°C, 19/14°C; 16h | Length of the ridge of the apical dome | Stereomicroscope | Fig.1 and Fig.2, pg. 58 |
| Bassiri et al. (1992)* | A632/W123 (Hybrid), Tp2l$^{+}$(Mutant) | Greenhouse; 24/20°C;16h | Length is half of the size of the apex | Stereomicroscope | Fig.1, pg.498 |
| Lejeune & Bernier (1996)* | B22 (Inbred) | Growth chamber; 24/18°C; 16h | Not stated | Binocular microscope | Fig.3, pg.220 |
| Fletcher et al. (2004)* | ‘Challenger’ sweet corn (Hybrid) | Field at Lincoln, Canterbury, New Zealand (2002); Five rates of P fertilisation | Not stated | Binocular microscope | Fig.4, pg.94 |
| Padilla & Otegui (2005)* | 33G26,34B15,34K77, 34N16 36G12 39A26 39R52, ANJOU 258, Monumental, DK312, DK315, DK615, DK664MG, DK682, DK752, DK834, B73xMo17 (Hybrids)<br>B73, Mo17 (Inbreds) | Field at University of Buenos Aires, Argentina; Five experiments (2002-2004) | Length is one third of the apex dome | Binocular microscope | Original data provided by authors |
| Andrieu et al. (2006)* | Dea (Hybrid) | Field at Grignon, France (2000); Two contrasting densities: 9.5 and 30.5 plants m$^{-2}$ | Length is above 25 µm | Binocular microscope | Fig.2, pg.1009 |
| Birch et al. (2007)* | C87, P3527 (Hybrids) | Field at Gatton, Australia; Two experiments (1999, 2000) | Length is above 25 µm | Stereomicroscope | Fig.7, pg.221 |
| Chenu et al. (2008)* | Dea (Hybrid) | Field at Mauguio, France; Three experiments (1995; 1997; 1998) | Length is above 50 µm | Stereomicroscope | Fig.1, pg.382 |
| Louarn et al. (2010)* | F286, F2 (Inbred) | Greenhouse; 24/20°C; Cold treatment (14°C/10°C) applied for one week; 16 hr | Length is half of the size of the apex | Binocular microscope | Fig. 2, pg.110 |
| Vidal et al. (2021)* | M52, M40 | Field at Grignon, France (2018) | Length is above 25 µm | Digital microscope | Fig. 3, pg.6 |

[1]Studies marked by the asterisk were selected for meta-analysis estimates.

**Table S2. Traits measured in platform and field experiments.**

| Trait Type | Trait | Units | Platform[1] | Field[1] |
|---|---|---|---|---|
| Canopy dimension | Blade length | cm | 154 (10) | 880 (114) |
| Canopy dimension | Blade width | cm | 154 (10) | 880 (114) |
| Canopy dimension | Sheath length | cm | 154 (10) | NA |
| Canopy dimension | Internode length | cm | 154 (10) | NA |
| Phenology | Collar appearance | Collar $d_{20^{\circ}C}^{-1}$ | 154 (10) | NA |
| Phenology | Leaf tip appearance | Leaf tip $d_{20^{\circ}C}^{-1}$ | 154 (10) | NA |
| Phenology | Total leaf number | Leaf rank | 154 (10) | 880 (114) |
| Phenology | Dominant ear rank | Leaf rank | 141 (10) | 880 (114) |

[1]Total number of plants measured, with number of genotypes indicated in paratheses.

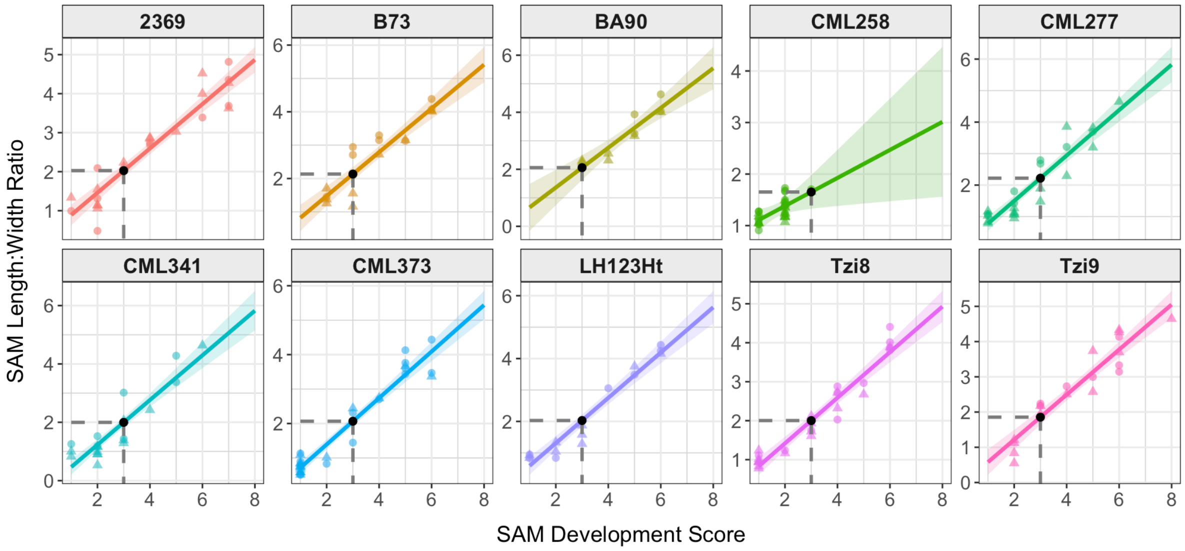


**Figure S1. Comparison between methods for determining tassel initiation of the shoot apical meristem.** Plots show results from the Night-Break platform experiment using a visual scoring system versus image-based quantification of shoot apical meristem (SAM) morphology. Data points correspond to the measurements from dissected plants, measured by two methods: the SAM length:width ratio estimated by image analysis (y-axis), and a developmental score assigned by visual classification (x-axis; a customized scoring scale [1-10] informed by Bonnett [1966] and Krüger [1984]). The Night-Break treatment had a significant but small effect on T.I. timing (approximately 1-2 leaf difference) but did not show a significant effect on the relationship between the scoring systems, so the data were pooled from both treatments. The fitted line and 95% confidence interval in each plot show the relationship from linear regression. The dashed line indicates that the qualitative classification score of 3, corresponding to T.I. according to Krüger (1984), maps to an average SAM length:width ratio of 2.0. This value was used to define T.I. with the image-based approach. Due to growth defects (buggy whipping) and late timing of T.I. in CML258, we excluded it from the main study but show the result here.

Bonnett, O. T. (1966). Inflorescences of maize, wheat, rye, barley, and oats: Their initiation and development. *Illinois Agricultural Experiment Station Bulletin*.

Krüger, G. H. J. (1984). The effect of photoperiod on the initiation of the staminate inflorescence in different hybrids of *Zea mays*. *South African Journal of Botany*, 3(1), 81–82. https://doi.org/10.1016/S0022-4618(16)30084-5

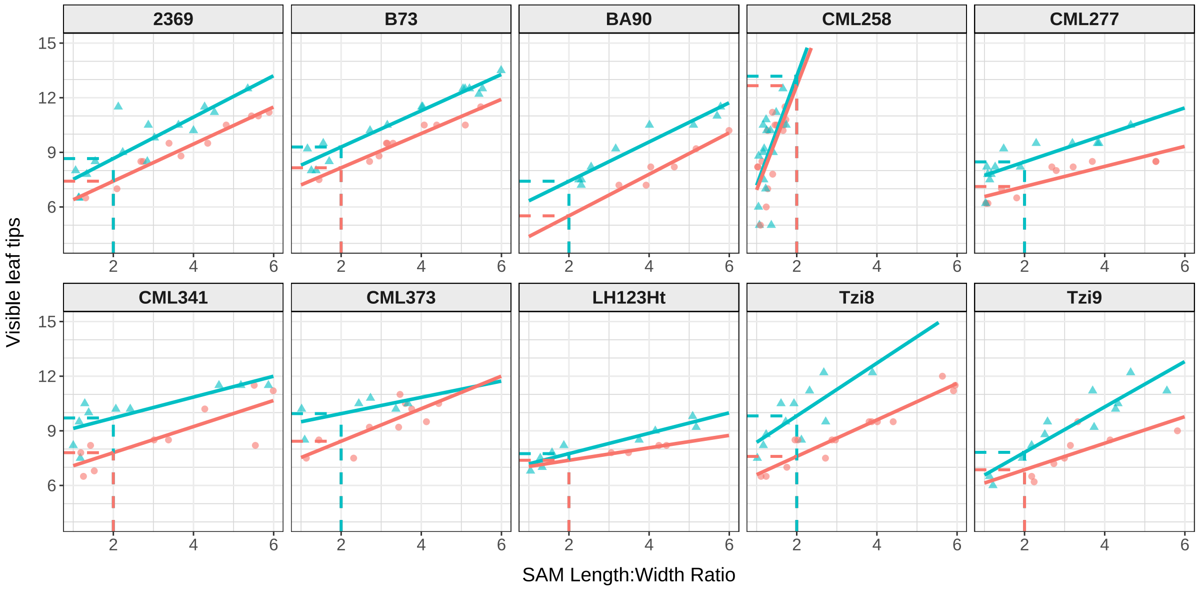


**Figure S2. Relationship between the visible leaf tip and development of the shoot apical meristem.** Plots show results from the Night-Break platform experiment used to empirically estimate T.I. timing. Data points correspond to the visible leaf tip number and the SAM length:width ratio for each serially dissected genotype tested in the two treatment conditions (blue=night-break; red=control). For each genotype × treatment combination, the linear regression between visible leaf tip number and SAM length:width ratio was used to estimate the leaf tip number at the fixed ratio of 2.0 (dashed lines), the ratio we determined to correspond with morphological shift to tassel initiation (Fig. S1). Due to growth defects (buggy whipping) and late timing of T.I. in CML258, we excluded it from the main study but show the result here.

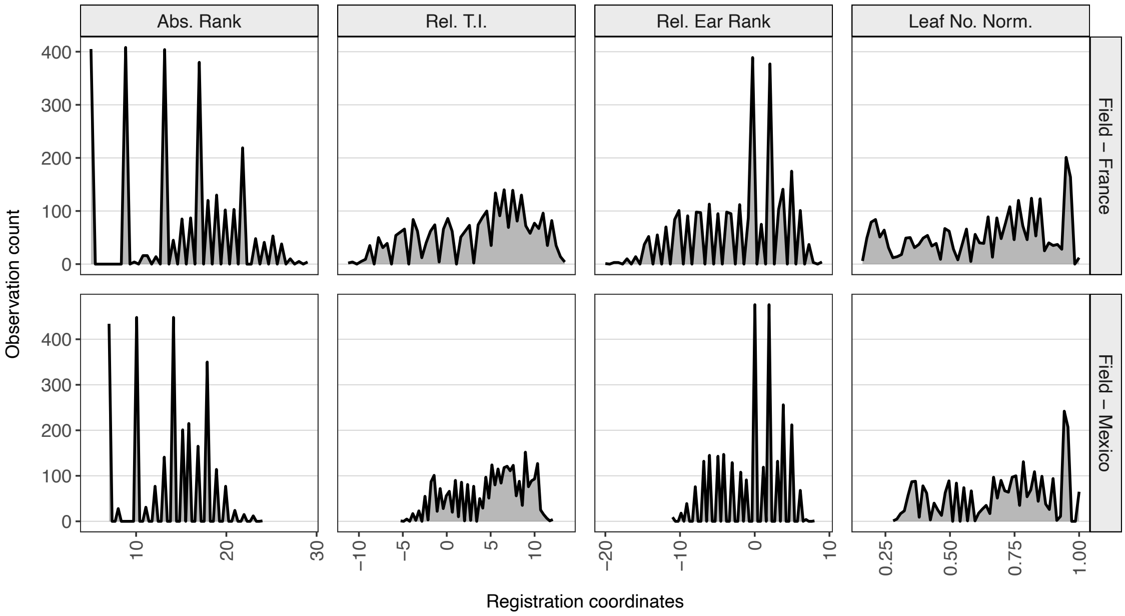


**Figure S3. Observation density for leaf blade length measurements from the field experiment.** Plots show the number of measurements across the coordinates of each registration system for the separate field environments (France and Mexico).

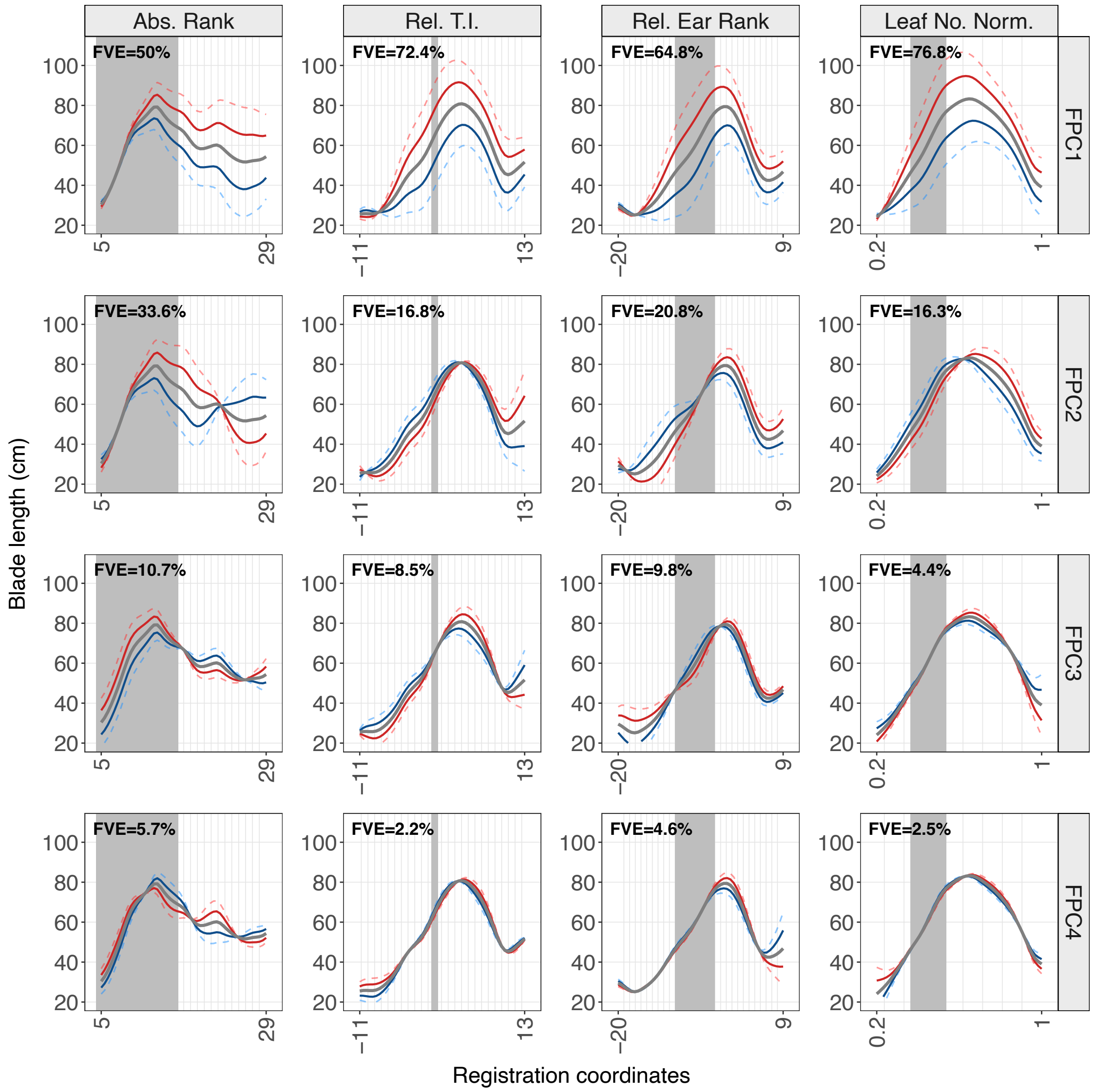


**Figure S4. Modes-of-variation for functional principal components by registration system.** The continuous grey line is the mean blade-length profile across environments. Coloured lines show synthetic profiles perturbed from the mean along each eigenfunction by one (solid) and two (dashed) standard deviations of the corresponding FPC score, in the positive (red) and negative (blue) directions. The percentage of variance explained by each FPC is indicated. Shaded regions show the range of tassel initiation across all plants included in the alignment.

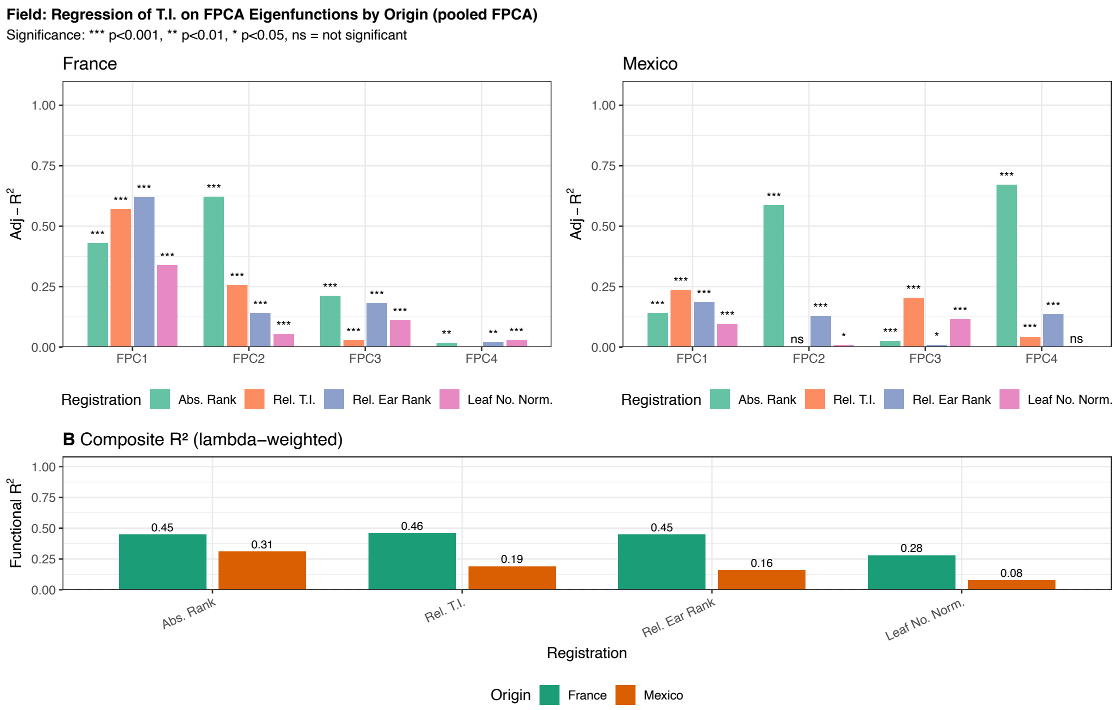


**Figure S5. Variation in functional principal components explained by tassel initiation.** After applying PACE to blade-length data pooled from both field environments, FPC scores were separated by environment and regressed on tassel initiation (T.I.). (**A**) Adjusted $R^2$ values from the regression analysis are shown for each FPC per environment (France and Mexico). (**B**) Composite $R^2$ values (see methods in the main text) quantify the total functional variation explained across FPCs per environment.
